# Abundant small RNAs in the reproductive tissues of the honey bee, *Apis mellifera*, are a plausible mechanism for epigenetic inheritance and parental manipulation of gene expression

**DOI:** 10.1101/2021.08.27.457896

**Authors:** Owen T. Watson, Gabriele Buchmann, Paul Young, Kitty Lo, Emily J. Remnant, Boris Yagound, Mitch Shambrook, Andrew F. Hill, Benjamin P. Oldroyd, Alyson Ashe

## Abstract

Polyandrous social insects such as the honey bee are prime candidates for parental manipulation of gene expression in offspring. Although there is good evidence for parent-of-origin effects in honey bees the epigenetic mechanisms that underlie these effects remain a mystery. Small RNA molecules such as miRNAs, piRNAs and siRNAs play important roles in transgenerational epigenetic inheritance and in the regulation of gene expression during development. Here we present the first characterisation of small RNAs present in honey bee reproductive tissues: ovaries, spermatheca, semen, fertilised and unfertilised eggs, and testes. We show that semen contains fewer piRNAs relative to eggs and ovaries, and that piRNAs and miRNAs which map antisense to genes involved in DNA regulation and developmental processes are differentially expressed between tissues. tRNA fragments are highly abundant in semen and have a similar profile to those seen in semen in other animals. Intriguingly we find abundant piRNAs that target the sex determination locus, suggesting that piRNAs may play a role in honey bee sex determination. We conclude that small RNAs play a fundamental role in honey bee gametogenesis and reproduction and provide a plausible mechanism for parent-of origin-effects on gene expression and reproductive physiology.

## Introduction

Parents can influence the phenotype of their offspring not only through the genetic contribution that they pass on but also through so-called ‘parental effects’. Maternal effects, the best-studied parental effect, occur when the mother influences the phenotype of the offspring, regardless of the offspring’s own genotype. For example, strong maternal effects can be caused by maternal deposition into oocytes of proteins or messenger RNAs (mRNAs) that are critical for early development of the zygote. If the mother is unable to provide these critical proteins/mRNAs, the resultant embryo dies regardless of its genotype. Maternal effects have been recognised by biologists for many decades whereas paternal effects are less well studied, partially due to the relatively small size of sperm and the commonly held belief that sperm contribute only their DNA to the fertilised egg. However, it is increasingly clear that paternal effects exist, although the mechanisms by which they are mediated is often unclear (Crean and Bonduriansky 2014).

Honey bees are a species in which parental manipulation of gene expression (such as via parental effects) is likely to evolve because i) females are polyandrous (mate with many males), ii) there is large investment in offspring colonies, the costs of which are borne by the parent colony collectively, but the majority of benefits accrue to a tiny number of offspring queens and their parents, iii) the value of male and female offspring to parents differs strongly (Haig 1999, 2000; Queller 2003; BURT and Trivers 2006; Haig 2010). Parental manipulation of offspring gene expression is particularly relevant for fathers, who die after mating (Winston 1987), and are therefore unable to directly enhance the reproductive success of their daughters. Indeed, a drone’s only opportunity to enhance the reproductive outcomes of his daughters is to use epigenetic manipulations that provide his daughters with an advantage over the daughters of other males (Haig 1992; Queller 2003; Drewell et al. 2012; Pegoraro et al. 2017). For example, a male might attempt to minimise additional matings by the queen that he has just mated with, potentially increasing the reproductive success of his own daughters (Liberti et al. 2019). Further, it has been repeatedly shown that larvae reared as queens comprise a non-random set of patrilines, suggesting that some males enhance the probability that their daughters will be reared as reproductive queens, potentially by epigenetic means (Osborne and Oldroyd 1999; Châline et al. 2002; Withrow and Tarpy 2018).

Small RNAs refer to non-coding RNA molecules between 18 and 50 nt long. Multiple classes of small RNAs are abundant in the semen of many animals and can alter offspring phenotypes when injected into zygotes (Vogel et al. 2010; Rassoulzadegan et al. 2006; Gapp et al. 2014; Grandjean et al. 2015; Benito et al. 2018; Rodgers et al. 2015; Chen et al. 2016a). Small RNAs are deposited maternally in *Drosophila melanogaster* (Brennecke et al. 2008; Akkouche et al. 2013) and recent evidence suggests that small RNAs are also deposited paternally and can influence gene expression in the next generation (Lempradl et al. 2021). Two examples demonstrate that biologically active small RNAs can be transferred from hymenopteran parents to offspring or from workers to larvae, providing a plausible mechanism by which epigenetic information could be transmitted between generations in honey bees. First, biologically active dsRNAs that confer acquired immunity are shared between honey bee generations via the glandular secretions that workers feed to larvae (Maori et al. 2019). Second, female jewel wasps (*Nasonia vitripennis* - of the same taxonomic order as honey bees) include RNA molecules in their eggs that determine the sex of their offspring (Verhulst et al. 2010).

Small RNAs are classified according to their size and function. The first-discovered small RNA molecules, collectively known as microRNAs (miRNAs), are short (c.a. 22 nt) non-coding RNAs that typically bind to the 3’ UTR region of messenger RNAs, causing translational inhibition and/or mRNA degradation (Cannell et al. 2008; Gebert and MacRae 2019). Maternally inherited miRNAs are involved in sex determination during embryogenesis in *Caenorhabditis elegans* (McJunkin 2018) and in mouse sperm, miRNAs degrade maternal mRNA stores in early zygotes to reprogram gene expression in the offspring (Rodgers et al. 2015). Some miRNAs are maternally deposited in *D*. *melanogaster*, where they play important roles in embryogenesis (Soni et al. 2013; Lee et al. 2014; Kugler et al. 2013a).

Piwi interacting RNAs (piRNAs) are a class of small RNA between 24-32 nt in size, that typically have a uridine at the 5’ end (5′U bias) (Gunawardane et al. 2007). piRNAs interact with PIWI proteins, a class of Argonaute protein, to repress transposable element (TE) activity in the germline of animals during meiosis (Girard et al. 2006), including in insects (Anand and Kai 2012; Brennecke et al. 2007). Additionally, piRNAs are involved in regulating gene expression in developing sperm cells and in somatic cells (Czech and Hannon 2016; Thomson and Lin 2009; Weick and Miska 2014). In *Drosophila* embryonic somatic cells, piRNAs destabilise/cleave target mRNAs to regulate embryonic development (Dufourt et al. 2017). In *Drosophila* germ cells piRNAs interact with Aubergine and a germline specific poly(A) polymerase to facilitate the localisation of essential germline mRNAs to the germ plasm, where they are protected and can be passed from generation to generation (Dufourt et al. 2017). Aubergine-piRNA-mediated epigenetic silencing of protein coding genes is well characterised in *D. melanogaster*, and can occur through piRNA induced silencing complexes (Rouget et al. 2010; Barckmann et al. 2015; Wang and Lin 2021) and/or spreading of piRNA-mediated heterochromatin into neighbouring loci (Lee 2015).

tRNA fragments (tRFs) are small RNA molecules that are derived from the cleavage of mature transfer RNAs (tRNAs) (Keam and Hutvagner 2015). The function of tRFs is mostly an enigma: some tRFs act similarly to miRNAs (Haussecker et al. 2010), while others interfere with global protein translation at the ribosome (Ivanov et al. 2011; Kim et al. 2017). Like miRNAs, tRF expression in *Drosophila* is age dependent. tRFs contain a seed region that has complementarity to 3’UTR regions of messenger RNAs and they interact with Argonaute proteins, suggesting that they form post-transcriptional RNA-induced silencing complexes (RISC) (Karaiskos et al. 2015) and/or are involved in influencing mRNA stability and transport (Göktaş et al. 2017). In mammalian sperm there is a global loss of piRNAs and an increase in tRFs and miRNAs that occurs during transit through the epididymis, a process that is essential for sperm maturation. tRFs have also been implicated in paternal transgenerational epigenetic inheritance (Sharma et al. 2016; Chen et al. 2016b; Cropley et al. 2016).

The central roles played by miRNAs, piRNAs and tRFs in spermatogenesis, nurturing of germ line cells, somatic gene regulation and epigenetic inheritance in other species make them prime mechanistic candidates for parental manipulation of offspring development in honey bees (He et al. 2009; Thomson and Lin 2009). Previous studies of small RNAs in *A. mellifera* have primarily focussed on ovary, embryo and thorax tissues to investigate caste determination, oviposition or phylogenetic similarities (Lewis et al. 2018; Pires et al. 2016; Chen et al. 2017; Wang et al. 2017). Here, we focus on small RNAs in reproductive tissues to investigate their potential roles in gametogenesis, parent-of-origin effects and epigenetic inheritance. We find that honey bee reproductive tissues have distinctive small RNA profiles, with ovaries and eggs having a higher proportion of piRNAs relative to other tissues, semen having a higher proportion of tRNA fragments, and spermatheca and testes having a higher proportion of miRNAs. Our miRNA and piRNA target prediction suggests that germ cells utilise small RNAs to regulate processes during development (post-fertilisation), and in gametogenesis and sex determination. Our findings place small RNAs front and centre as the mechanism that mediates the parent-of-origin effects that are phenotypically observed in honey bees, but which cannot be ascribed to other epigenetic mechanisms, notably DNA methylation (Remnant et al. 2016; Guzman-Novoa et al. 2005; Wu et al. 2020; Galbraith et al. 2016; Kocher et al. 2015; Oldroyd and Yagound 2021).

## Results

### Length distribution and biotype analysis of small RNAs

We surveyed the small RNA populations in the reproductive and germline tissues of Australian commercial honey bees (mainly *A. m. ligustica* heritage). We extracted RNA and generated small RNA libraries from a range of male (testes and semen) and female (spermatheca and ovary) reproductive tissues, as well as eggs from mated and virgin queens. The spermatheca is the organ in which sperm are stored, which a queen uses to fertilise female-destined eggs throughout her life. Sequencing of these libraries generated 12 to 35 Mb of reads per sample. On average 65% of reads mapped to the *A. mellifera* 4.5 genome. We first classified the small RNAs by size and biotype. Strikingly, the RNA populations varied greatly in size and biotype depending on their tissue of origin (Figure 1A, Supplemental Figure 1).

**Figure 1.**
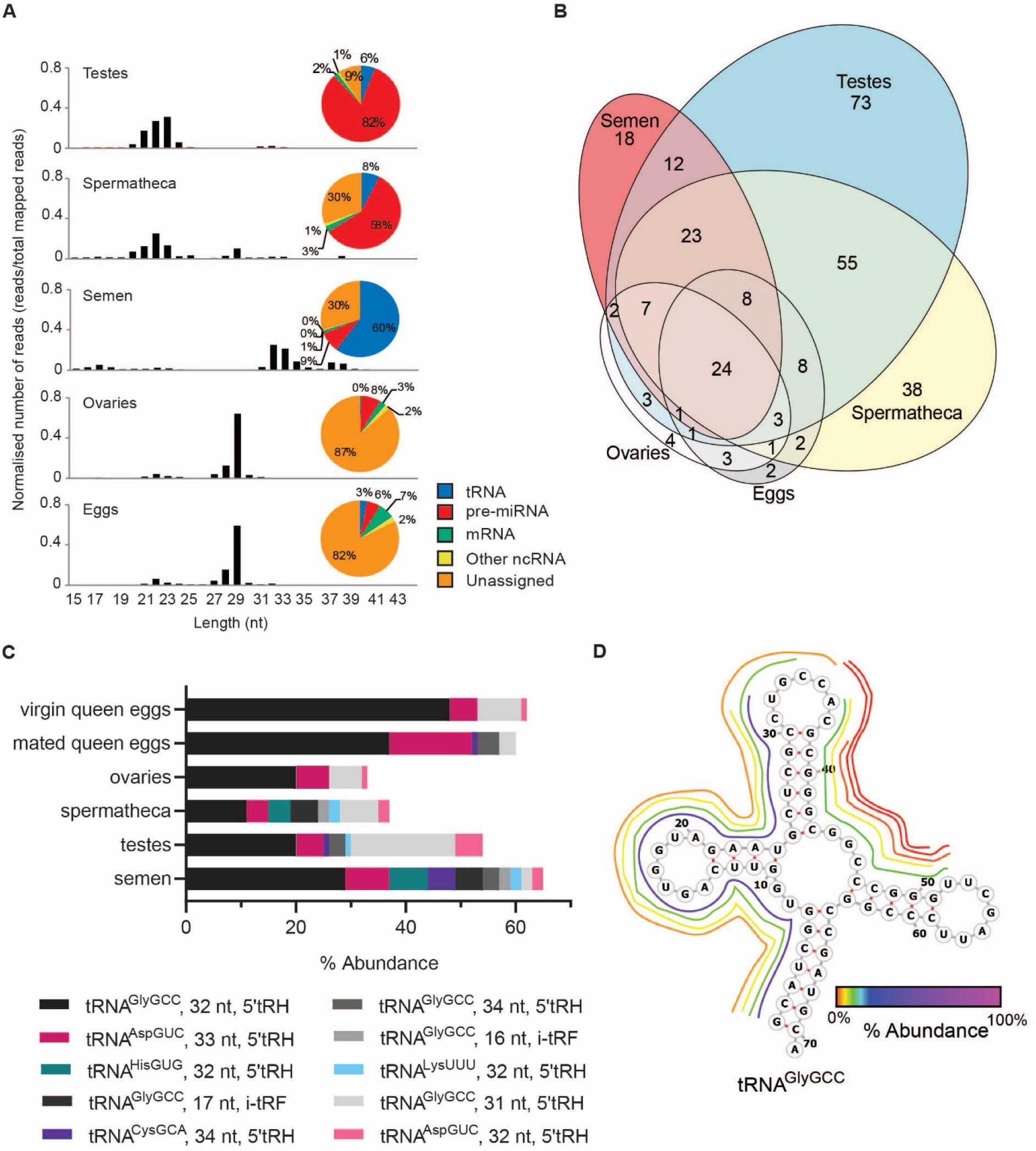
Small RNA species in the reproductive tissues of the honey bee. **A**. Size distribution of small RNAs between 13 and 43 nt in length mapped to the Amel 4.5 honey bee genome. Pie charts show the proportion of mapped reads that align to annotations of each biotype (tRNA, pre-miRNA, mRNA, other ncRNA and unannotated) for each tissue type. **B**. Euler plot illustrating number of novel miRNAs discovered in each reproductive tissue type and the overlap between tissue types. **C**. The 10 most abundant tRFs in semen and their relative abundance in other tissue types. tRF isoforms that originate from the same tRNA are shown as different shades of the same colour. **D**. A representation of the abundance of tRNA fragments derived from the tRNA^GlyGCC^ in semen. The origin of each tRF is shown by its position around the molecule and relative abundance is indicated by colour.

Mapped reads from spermatheca and testis samples had a prominent peak at 21-23 nt which is the expected size of miRNAs. 58% and 82% respectively mapped to known precursor miRNAs from miRbase (Figure 1A). In addition to honey bee miRNAs contained in miRbase, we identified 301 novel miRNAs using miRDeep2 (Friedländer et al. 2012). The majority of these novel miRNAs were identified in the testes and spermathecal libraries (Figure 1B) (Supplemental Figure 1B, Supplemental Table S1). StemLoop qPCR (Hurley et al. 2011) was conducted to validate three of the novel miRNAs that were differentially expressed between tissue types, according to small RNA sequencing. In all three cases the StemLoop qPCR results mirrored the expression levels suggested by the small RNA sequencing (Supplemental Figure 1C).

In contrast to all other samples, semen samples had an abundance of mapped reads in the 32-33 nt range. Biotype analysis revealed that 60% of these reads mapped to sub-regions of tRNAs and were therefore tRNA fragments (tRFs) (Figure 1A). Following the classifications of Loher and colleagues (Loher et al. 2017), tRFs were classified as being either 5′ or 3′ tRNA halves (tRH), 5′ or 3′ tRFs or internal tRFs (itRFs). Figure 1C shows the top 10 tRFs in semen, and their proportional abundance in other tissues. A full list of the relative abundance of tRFs across all tissue types is provided in Supplemental Table S2. The levels of individual tRFs appear to be highly regulated because their abundance differed greatly among tissues. For example, although the most abundant isodecoder, tRNA^GlyGCC^, generated 30-60% of all tRFs per tissue type, the first and second most abundant tRNA^GlyGCC^ itRFs in semen were present at much higher levels in semen and spermatheca than in all other tissues (Figure 1C). This difference indicates that tRF production is tissue-specific, and not simply a consequence of tRNA degradation. Strikingly, the high abundance of tRFs in semen was not observed in the spermatheca (Figure 1A). This suggests that tRFs present in semen are carried in the seminal fluid and not by the spermatozoa, and/or that female contribution of small RNAs to the spermathecal fluid is abundant.

Mapped reads from egg and ovary samples showed a peak at 29 nt. This peak was also evident in the spermathecal samples but was relatively smaller than the larger peak at 21-23 nt (Figure 1A). The high abundance of 29nt length reads was evident in both mapped and unmapped reads (Supplemental Figure 1B), suggesting that many were from repetitive regions that were not placed in the Amel 4.5 genome assembly. Egg and ovary samples also had a large proportion of total reads (~85%) that did not map to any recognised category of small RNA. The 29 nt reads are within the size range of piRNAs and nucleotide frequency plots generated using 26-31 nt reads showed a strong bias towards 1U in ovaries and eggs (Supplemental Figure 2). We therefore hypothesised that these 29 nt small RNAs were piRNAs. To generate a list of putative piRNAs for each tissue, reads mapping to other known ncRNAs (i.e. tRNAs, rRNAs, miRNAs) and reads below 26nt or above 31nt were removed.

### piRNA cluster analysis

piRNAs are often transcribed from pericentromeric and telomeric heterochromatic regions, termed piRNA clusters, that are characterized by an abundance of transposable element (TE) remnants (Andersen et al. 2017). Using three algorithms: a custom script (CREST), proTRAC v2.4.2, and piClust, we identified 199 putative honey bee piRNA clusters totaling 3,113.3 kB (1.25% of the genome) (Supplemental Table S3). Of the 199 clusters, 113 (57%) were located on unmapped scaffolds, probably due to the association with repetitive sequences that are hard to map (Huang et al. 2017).

In contrast to *D. melanogaster* (Czech and Hannon 2016; Akkouche et al. 2017), we found that most honey bee piRNA clusters (171 of 199) are uni-directional and equally present on both genomic strands (Supplemental Table S3). This supports the suggestion of Wang et al that species other than *Drosophilids* are incapable of dual-stranded piRNA cluster activity (Wang et al. 2017).

Clusters ranged in size from 1kB (Cluster 28) to 152 kB (Cluster 98) (Supplemental Table S3). Of the 199 clusters, 69 were identified in only one tissue, whereas only eight clusters were identified in all five tissues (Figure 2A), demonstrating strong tissue-specific regulation of piRNA expression. The vast majority of putative piRNA reads mapped to clusters for ovaries (99%), eggs (99%), and spermatheca (90%). In contrast, only 56% of putative piRNAs mapped to clusters for testes and 23% for semen, suggesting that many of the piRNA-sized reads in these two tissues were not *bona fide* piRNAs, or are produced non-canonically. Of the piRNA reads that mapped to clusters, over 50% mapped to cluster 83 in eggs, ovaries and spermatheca (Figure 2B). Cluster 83 contains many transposable element remnants (Figure 2C) (predominantly large retrotransposon derivatives (LARDs)) and is the main piRNA-generating cluster in female tissues. This shows that honey bee piRNA clusters, like those in *Drosophila* (Brennecke et al. 2007), resemble transposon graveyards. In semen, 75% of clustered piRNA reads map to cluster 8, which contains one large peak of reads but does not overlap any TE or other genomic feature. This again suggests that TEs are not associated with piRNA-sized reads in semen.

**Figure 2.**
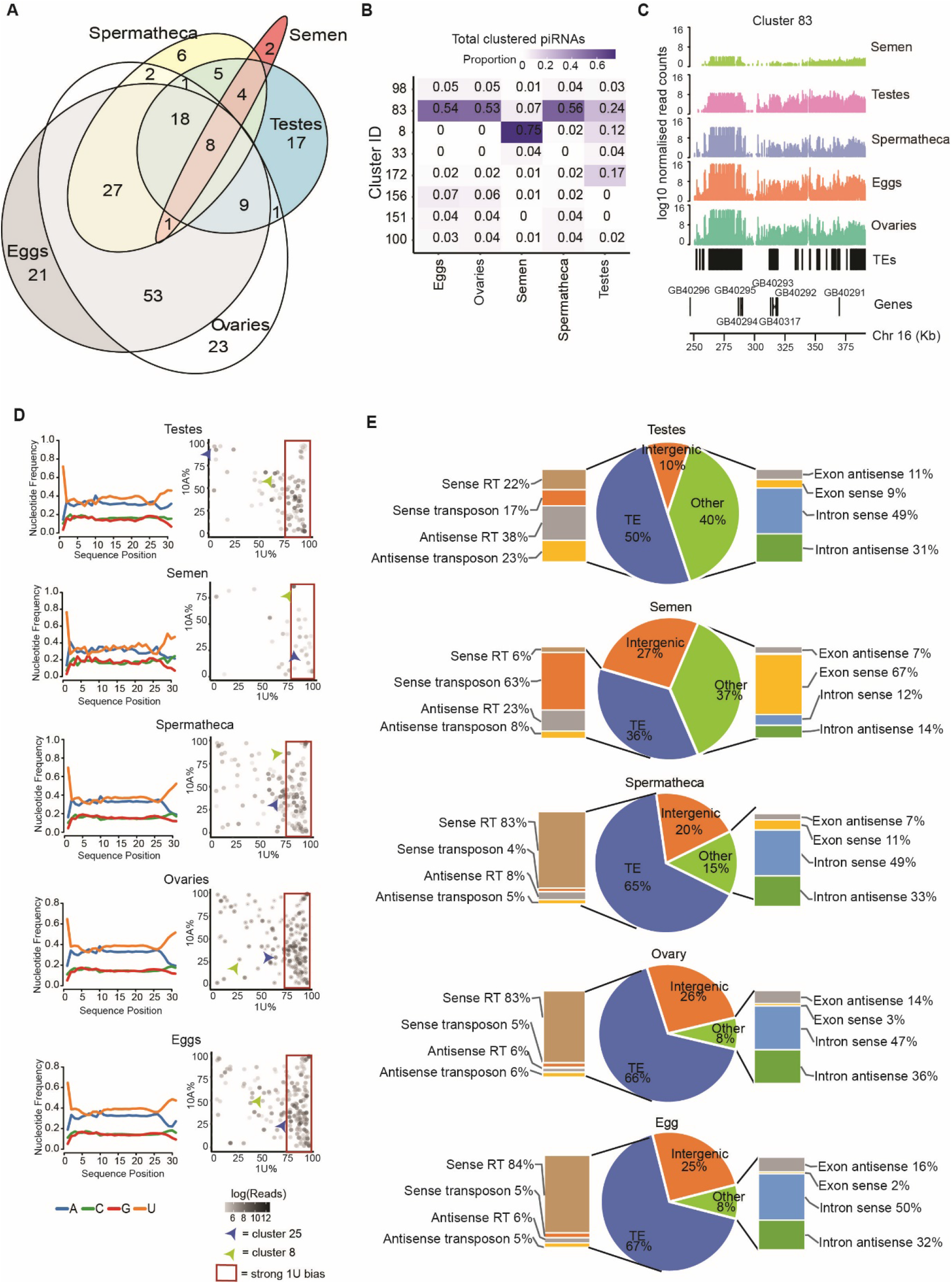
Analysis of piRNAs in the reproductive tissues of the honey bee. **A**. Euler diagram showing the overlap of clusters identified within each individual tissue. **B**. Table of the proportion of clustered total/uncollapsed piRNAs that align to the top three most active piRNA clusters for each tissue. **C**. Coverage of piRNAs at Cluster 83 for each tissue. Y-axis values are log10 RPM (reads per million) normalised values of total reads. **D**. Left: Nucleotide frequency plots for unique piRNAs in ovaries, eggs, testes, semen and spermatheca samples. Right: The 1U and 10A bias of each piRNA cluster, detected within each tissue individually. Red box indicates clusters with a 1U bias. Cluster 25 (blue arrow) and cluster 8 (green arrow) are examples of the same piRNA cluster with different 1U and 10A frequencies between tissue types. **E**. Feature mapping of total piRNAs sense and antisense to retrotransposons and DNA transposons (left) as well as introns and exons (right)

Previous analysis of whole tissue honey bee larvae suggested that honey bees utilise ping pong biogenesis in the production of piRNAs (Wang et al. 2017). To determine if this holds true for reproductive tissues, we assessed two signatures of ping pong biogenesis (Czech and Hannon 2016): the prevalence of a 1U/10A nucleotide bias and complementary sequence overlaps spanning 10 bp. Nucleotide frequency analysis of clustered piRNAs (in contrast to all piRNA-length reads) showed a strong 1U bias in all tissues but no strong 10A bias (Figure 2D), indicating that primary piRNAs are the most common overall. However, when 1U/10A ratios were plotted for each individual cluster it was apparent that some clusters had both a 1U and 10A bias, and that the extent of this bias differed between tissues (Figure 2D). Furthermore, a 10 bp overlap between reads was enriched in testes and spermatheca, present in ovaries and eggs, but conspicuously absent in semen (Supplemental Figure 3A). This suggests that some ping pong cycling is present in tissues except semen. We also measured piRNA phasing by plotting the 3’-5’ end distances between adjacent piRNAs. In *D. melanogaster* primary piRNA biogenesis PIWI proteins use piRNAs as guides to fragment a pre-piRNA into a string of tail-to-head phased pre-piRNAs that are further processed into mature piRNAs (Gainetdinov et al. 2018). We detected a significant signature of phased piRNA in semen (Z = 3.38, p < 0.001), but not in other tissues (Supplemental Figure 3B). This indicates that primary piRNAs dominate in semen.

Overall, these data support previous findings that honey bees have ping-pong cycling capability (Wang et al. 2017), but strongly suggest that this activity is highly locus-specific, and not a general feature of piRNAs biosynthesis. piRNAs that are present in semen are mostly primary piRNAs – unsurprising, given that spermatozoa are mostly transcriptionally inert (see Discussion).

### piRNA targeting

Most piRNA-length reads in maternal tissues mapped to TE-dense piRNA clusters. We hypothesized that semen and testes would have fewer TE-associated piRNA-length reads, as fewer piRNA clusters were identified in these tissues. As expected, we found that less than half of piRNA-length reads mapped to TEs in semen and testes, compared to around 65% in maternal tissues (Figure 2E). Furthermore, TE-mapping piRNAs in ovaries, eggs and spermatheca predominantly mapped to retrotransposons in the sense direction, whereas piRNAs in semen predominantly mapped to DNA transposons (Figure 2E), revealing clear tissue specificity in the types of TE targeted by piRNAs. Of the TEs that have been confidently assigned to TE families (more recently active TEs) (Elsik et al. 2014), the class II transposon Mariner/TC1 was highly targeted by piRNAs in all tissues except semen, in which the LINE retrotransposon R2 was the most targeted (Supplemental Table S4). Of putative piRNA reads that map to genes, most align sense to introns for all tissues except semen, in which the majority align sense to exons (Figure 2E). The relative lack of piRNAs in semen and the stark difference of targeting and abundance of piRNA-length reads in paternal tissues compared to all other tissues suggests that piRNAs are a mechanism by which maternal effects could be mediated.

### Differential expression and target analysis

Principal component analysis (PCA) revealed that miRNA, piRNA and tRF profiles differed between tissue types (Figure 3A). To identify small RNAs that were contributing to the differences between tissues we performed pairwise differential small RNA expression analysis for each tissue (Figure 3B, Supplemental Table 3).

**Figure 3.**
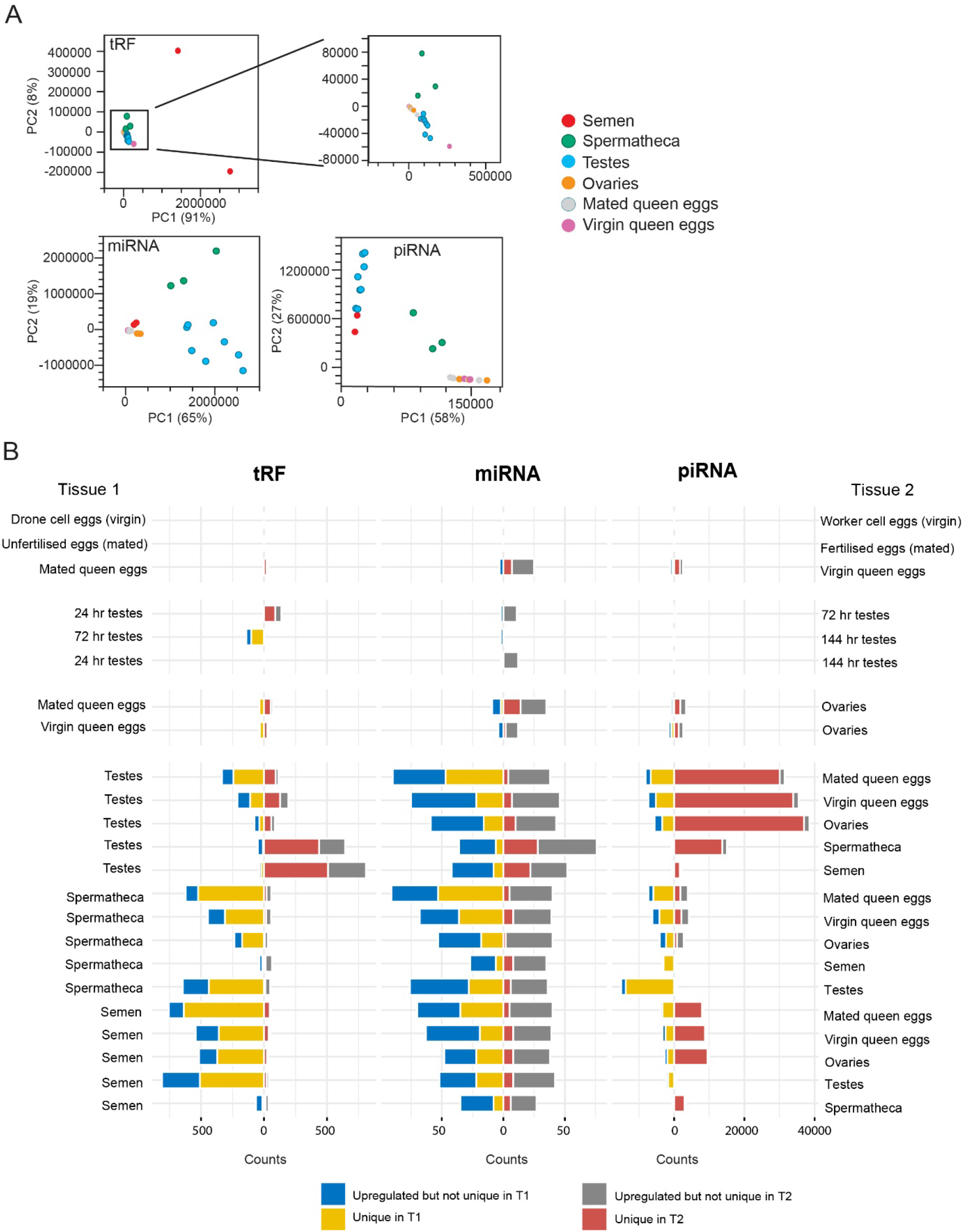
Differential expression analysis. **A**. Principal component analysis (PCA) plots on tRF (top), miRNA (bottom left) and piRNA (bottom right) profiles between tissue types. **B**. Differential tRF, miRNA and piRNA expression for pairwise tissue comparisons. Counts of differentially expressed and/or uniquely expressed small RNAs are on the x-axes. Red and grey bars extending to the right refer to the number of small RNAs that are uniquely (red) expressed or upregulated but not unique (grey) in tissue 2 relative to tissue 1. Conversely, the yellow bar extending leftward from the axis refers to the count of uniquely expressed small RNAs expressed in tissue 1 relative to tissue 2, the blue bar refers to the count of upregulated but not uniquely expressed sRNAs.

#### Ovaries and eggs

There is preliminary evidence that queens may use supplementary epigenetic mechanisms to influence the sex of their eggs (Oldroyd et al. 2008). However, we found no difference in the abundance of small RNAs between eggs from mated queens collected from worker cells (fertilized) and eggs collected from drone cells (unfertilized) (Figure 3B, row 2), indicating that at this age (<24hrs old) there is no difference in small RNA profiles between fertilised and unfertilised eggs. We therefore combined the data sets from mated queen eggs obtained from drone cells and worker cells into a single data set (mated queen eggs). We then compared sRNA profiles between eggs laid by virgin queens in worker cells with eggs laid in drone cells (both unfertilised). While we did not detect any differentially expressed (DE) miRNAs or tRFs we did detect 107 DE piRNAs (out of ~2.5 million piRNAs) (Figure 3B, row 1). Given the minimal differences between these two classes of virgin queen eggs we also pooled these datasets together (virgin queen eggs) for greater statistical power for comparisons between mated queen eggs and virgin queen eggs. We identified 39 tRFs, 30 miRNAs and 3,586 piRNAs that were DE between eggs from mated queens and eggs from virgin queens. The majority of these were upregulated in eggs laid by virgin queens (Figure 3B, Figure 4A). Due to these differences, we kept the virgin and mated queen egg datasets separate for comparisons with other reproductive tissues.

**Figure 4.**
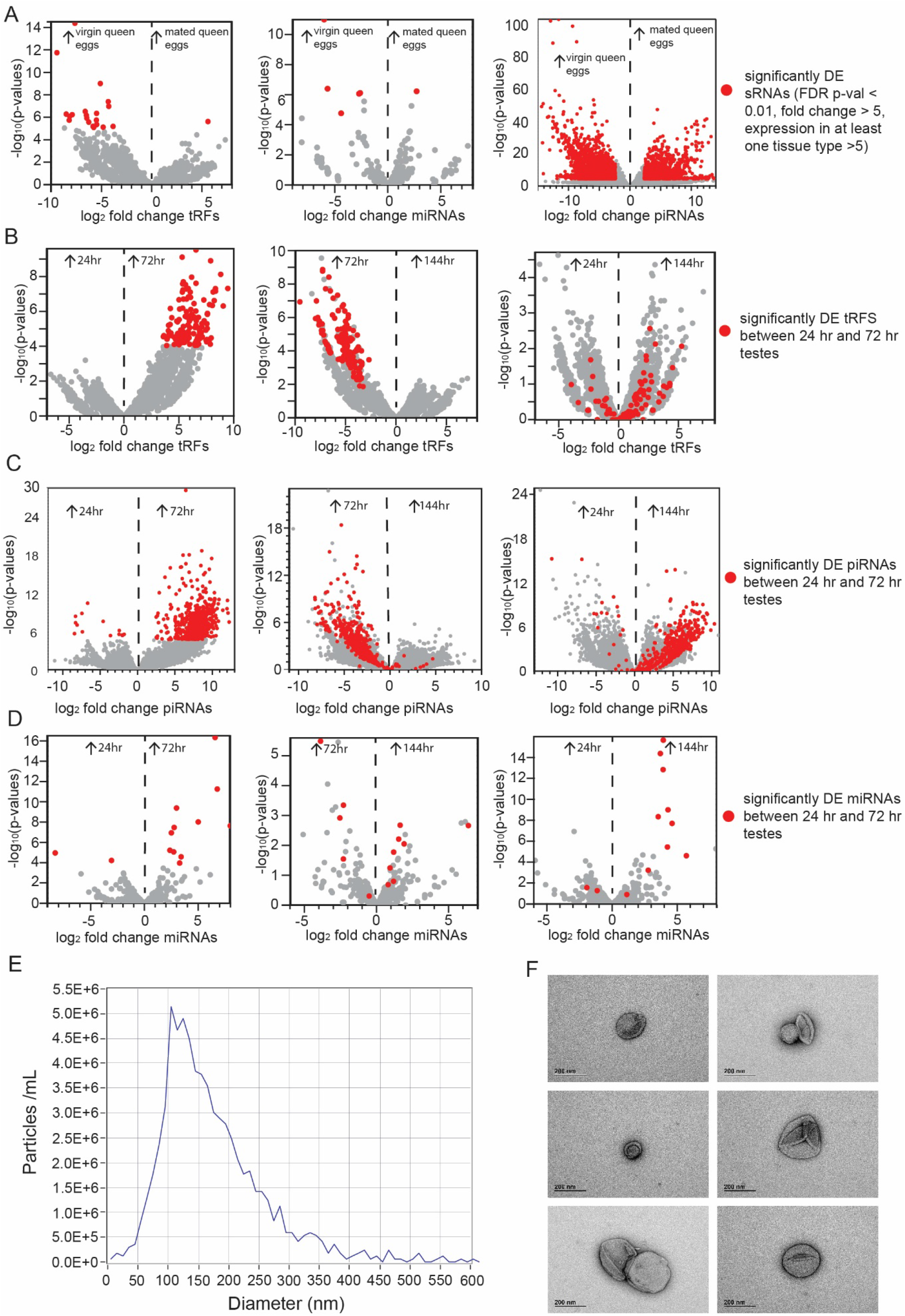
**A**. Volcano plots indicate the patterns of significantly (FDR-corrected) upregulated tRFs (left), miRNA (middle) and piRNAs (right) between eggs laid from virgin queens and eggs laid from mated queens. Arrows refer to upregulation in either virgin or mated queen eggs. **B**. **C. and D**. Volcano plots showing differentially expressed tRFs (B), piRNAs (C) and miRNAs (D) between testes from 24 hr pupae and 72 hr pupae (left), 72 and 144 hr (middle) and 24 and 144 hr (right). Red dots in all panels indicate the upregulated small RNAs from the 24-72 hr comparisons, grey dots are all small RNAs analysed. **E**. Zetaview measurement of particles present in honey bee semen indicating particle size (nm) and concentration (particles/mL). **F**. Representative transmission electron micrographs of potential EVs in honey bee semen.

The small RNA expression profiles in the eggs derived from both virgin and mated queen eggs were more similar to those seen in ovaries than to other tissues. This is expected, as the ovaries are the site of oogenesis and egg maturation, and the ovaries that we collected came from laying queens containing hundreds of developing eggs. Nonetheless, differentially expressed tRFs, miRNAs, and piRNAs were observed between ovary and egg samples (Figure 3B, Supplemental Table S5). Differentially expressed small RNAs, particularly miRNAs and piRNAs, tended to be upregulated in ovaries. Any differentially expressed small RNAs must originate in either the maternal ovarian tissue, arise from differential transport of miRNAs into mature eggs, or arise due to differential expression early in egg development. The small RNAs upregulated in the ovaries are likely to be derived from the maternal ovarian tissue and not from the developing oocytes, while the small RNAs upregulated in the eggs must come from either active transport into the developing oocyte, or early expression in the embryo. Of the seven miRNAs upregulated in ovaries (relative to both mated and virgin eggs) three (mir-989, mir-278 and mir-12) have been implicated in ovary development or oogenesis in insects (Kugler et al. 2013b; Song et al. 2018; Macedo et al. 2016).

#### Developing Testes

During spermatogenesis meiosis occurs as pupation commences and ends with the appearance of clustered spermatids in red-eyed pupae (Lago et al. 2020). To investigate whether paternally deposited small RNAs are produced during this process, we compared the testes samples from three different stages of pupal development; pink-eyed (24 hrs from the commencement of pupation), red-eyed (72 hrs) and brown-eyed (144 hrs). There were 139 differentially expressed tRFs between the developing testes of pink-eyed and red-eyed pupae and 140 differentially expressed tRFs between red-eyed and brown-eyed testes (Figure 4B, Supplemental Table S5). Strikingly, all differentially expressed tRFs were upregulated in the testes of red-eyed pupae, for each comparison. However, no tRFs were differentially expressed between pink-eyed and brown eyed pupae (Fig 4B). These results imply a peak of tRF expression at around 72 hrs of pupal development with subsequent downregulation. Like tRF expression, there is a drastic increase in piRNA expression within the testes between 24 and 72 hrs of pupal development (Figure 4C). These piRNAs show reduced abundance between 72 and 144 hrs, but not as strongly as tRFs. In contrast, miRNAs that are upregulated at 72 hr testes relative to 24 hr testes tend to remain upregulated in 72 hr testes and 144 hr testes, representing a trend towards increased miRNA production as the testis develop (Fig 4D). Of tRFs that were significantly upregulated at 72 hours (red-eyed pupae) relative to other pupal stages, many multi-mapped to eleven genes, two of which are important gametogenesis genes in *D. melanogaster*; *Hephaestus (heph)* (Robida et al. 2010; Sridharan et al. 2016) and *CUGBP Elav-like family member 2 (bru-2)* (Dasgupta and Ladd 2012). 72 hours post-fertilisation (red-eyed pupae) coincides with the end of meiosis and the start of spermiogenesis (Lago et al. 2020): the striking changes in small RNA expression during this window strongly suggest that small RNAs play an important role in spermatogenesis. Might they also influence gene expression in the next generation? For further comparisons we pooled all testes samples together.

#### Spermatheca, semen and testes

In order to determine whether paternal small RNAs are present in tissues relevant to the production of the next generation we considered the three environments encountered by honey bee sperm prior to fertilisation: the pupal testes, the adult seminal vesicles and the period of storage in the spermatheca (Bishop 1920a, 1920b). Strikingly, although tRFs made up 60% of small RNAs in semen but only 8% of spermathecal small RNAs by abundance (Figure 1B), there are fewer DE tRFs between semen and spermatheca samples than between other tissue types, suggesting high similarity between semen and the semen-storage organ (Figure 3B). In contrast, the stark difference in tRF abundance between semen and testes suggests that a large component of the tRF content in semendoes not come from the testes but is a component of the seminal fluid, which is produced in the accessory glands (Snodgrass 1910). This observation accords with the previous observation that tRFs are upregulated in the testes of red-eyed pupae but subsequently downregulated in brown-eyed pupae. The similarity in tRF expression between semen and spermatheca suggests that tRFs are a small RNA molecule uniquely placed to provide a paternal influence on early embryonic development.

Small RNAs are often present in extracellular vesicles (EVs) which have been shown to be present in semen of various organisms (Vojtech et al. 2014; Zhao et al. 2020; Chan et al. 2020). We performed particle tracking and electron microscopy of freshly collected honey bee semen and detected particles of the correct size and shape to be EVs (Figure 4E, F). If EVs are indeed present in honey bee semen they are a plausible mechanism by which small RNAs could be transmitted paternally.

A different trend was observed for piRNAs between semen, spermatheca and testes. There were few DE piRNAs between semen and testes. In contrast, the spermatheca was strongly different, with approximately 28-fold higher expression of unique piRNAs than in testes (Figure 3B, Supplemental Table S5). The abundance of piRNAs in the spermatheca is most likely due to piRNA expression in the maternal cells of the spermatheca itself, or in the maternal component of the spermathecal fluid, but probably not from spermatozoa.

The miRNA compositions between semen, spermatheca and testes are different again. There are few miRNAs that are unique to the testes, while there are many uniquely expressed miRNAs in the semen and spermatheca. Although miRNAs make up 82% of small RNAs in the testes, and 58% in the spermatheca, they comprise only 9% of small RNAs in semen. There are slightly fewer miRNAs upregulated in semen relative to spermatheca. This could suggest that there is a miRNA component to the seminal fluid (although substantially less than the tRF contribution shown above), and a maternal miRNA contribution to the spermathecal fluid. The seven most upregulated miRNAs in spermatheca relative to semen are also more abundant in spermatheca relative to all other tissues but are of conspicuously low abundance in eggs (Supplemental Figure 4). This suggests that these miRNAs regulate genes important to early embryogenesis. Indeed mir-317, mir-277 and mir-278 all play roles in insect development (Shen et al. 2020; Yang et al. 2016; Song et al. 2018).

#### Female tissues compared to male tissues

We sought to investigate how small RNAs differ between male and female reproductive tissues, to gain insight into how they may utilise different strategies to regulate gene expression and genomic stability or contribute small RNAs to the next generation. Semen and spermatheca express many more tRFs than do testes, ovaries, and eggs (Figure 3B, Supplemental Table S5), and thus tRFs are prime candidates for epigenetic marks passed by fathers to offspring. However, at our significance thresholds we did not detect any tRFs present in semen, spermatheca and fertilised eggs from mated queens that were not also in unfertilised eggs from mated queens.

Overall, paternal tissues and spermatheca have more abundant and uniquely-expressed miRNAs relative to eggs and ovaries (Figure 3B, Supplemental Table S5), although not to the same degree as tRFs. GO analysis of miRNA target genes showed that DNA binding activity involved in transcription regulation was often enriched for both tissues of a comparison, suggesting that miRNAs that are differentially expressed between tissues might act on/suppress different DNA-transcription machinery to regulate the activity of large cellular pathways (Supplemental Table S5). In contrast to tRFs and miRNAs, many more piRNA species are strongly upregulated in maternal tissues relative to testes and semen (Figure 3B, Supplemental Table S5). This is further evidence that piRNAs are expressed in the maternal reproductive tissues and suppressed in the paternal reproductive tissues. In addition to mapping to TEs, piRNAs also map to genes. GO-term analysis of genes targeted by piRNAs in ovaries and eggs relative to semen and testes (and presumably therefore downregulated in maternal tissues) were enriched for terms related to glutamate receptor signalling. Conversely, genes targeted by piRNAs in semen and testes relative to eggs and ovaries were enriched for terms related to transcription and development. Genes/clusters which have notably different antisense-piRNA targeting between tissue types are listed in Table 1, and the top 10 piRNA-targeted genes for each tissue pairwise comparison are listed in Supplemental Table S6. Full GSEA output is in Supplemental Table S7. The targeting of genes by piRNAs suggests that piRNAs also regulate gene expression and developmental pathways.

**Table 1.** Notable piRNA-targeted genes and clusters.

| piRNAs<br>upregulated in | Relative<br>to | Gene ID<br>(Beebase/Re<br>fseq) | Gene<br>Name | Gene function |
| --- | --- | --- | --- | --- |
| Eggs, ovaries and spermatheca | Semen and testes | GB55060/LOC724287 | *rhomboid related protein 3-like* | The rhomboid superfamily of intramembrane serine proteases are involved in developmental patterning as well as mitochondrial fusion during spermatogenesis (McQuibban et al. 2006; Bier et al. 1990). |
| Semen | Testes, eggs and ovaries | GB45868/LOC413081 | *F-box/WD repeat-containing protein 7* | This gene is a homolog of *archipelago* in *Drosophila*, required for endocycles and mediates the negative regulation of cell growth by protein ubiquitination (Moberg et al. 2004; Shcherbata et al. 2004) |
| Semen | Spermatheca, ovaries, testes and eggs | GB54783/LOC408402 | *Acetyl-coenzyme A transporter 1* | This gene is implicated in the developmentally regulated modification of O-acetylation of gangliosides in mammals (Kanamori et al. 1997) |
| Ovaries, eggs and spermatheca | Semen and Testes | Cluster 20 | Sex determination locus | Genes implicated in sex determination in *Apis mellifera* including the *complementary sex determiner* and *feminizer* genes. |

Intriguingly, piRNA cluster 20 is much more active in ovaries, eggs and spermatheca relative to testes and semen (Table 1). Cluster 20 is a unidirectional cluster that maps antisense to most of the genes located in the cluster. Reads map along the length of the cluster and although there are transposable elements present in the cluster, they are not exclusively targeted. Cluster 20 overlaps the *complementary sex determiner* gene (*csd*), *feminizer* gene (*fem)* and GB47023/LOC408733 (*pinocchio*) (Figure 5) (Beye et al. 2003). *csd*, situated within the sex determination region on chromosome 3, determines the sex of the bee and arose from duplication of the *feminizer* gene (Hasselmann et al. 2008). Bees heterozygous for *csd* develop into females, whereas bees hemizygous at this locus develop as haploid males (Beye et al. 2003). The high number of piRNAs at this locus suggest a role for piRNAs in sex determination.

**Figure 5.**
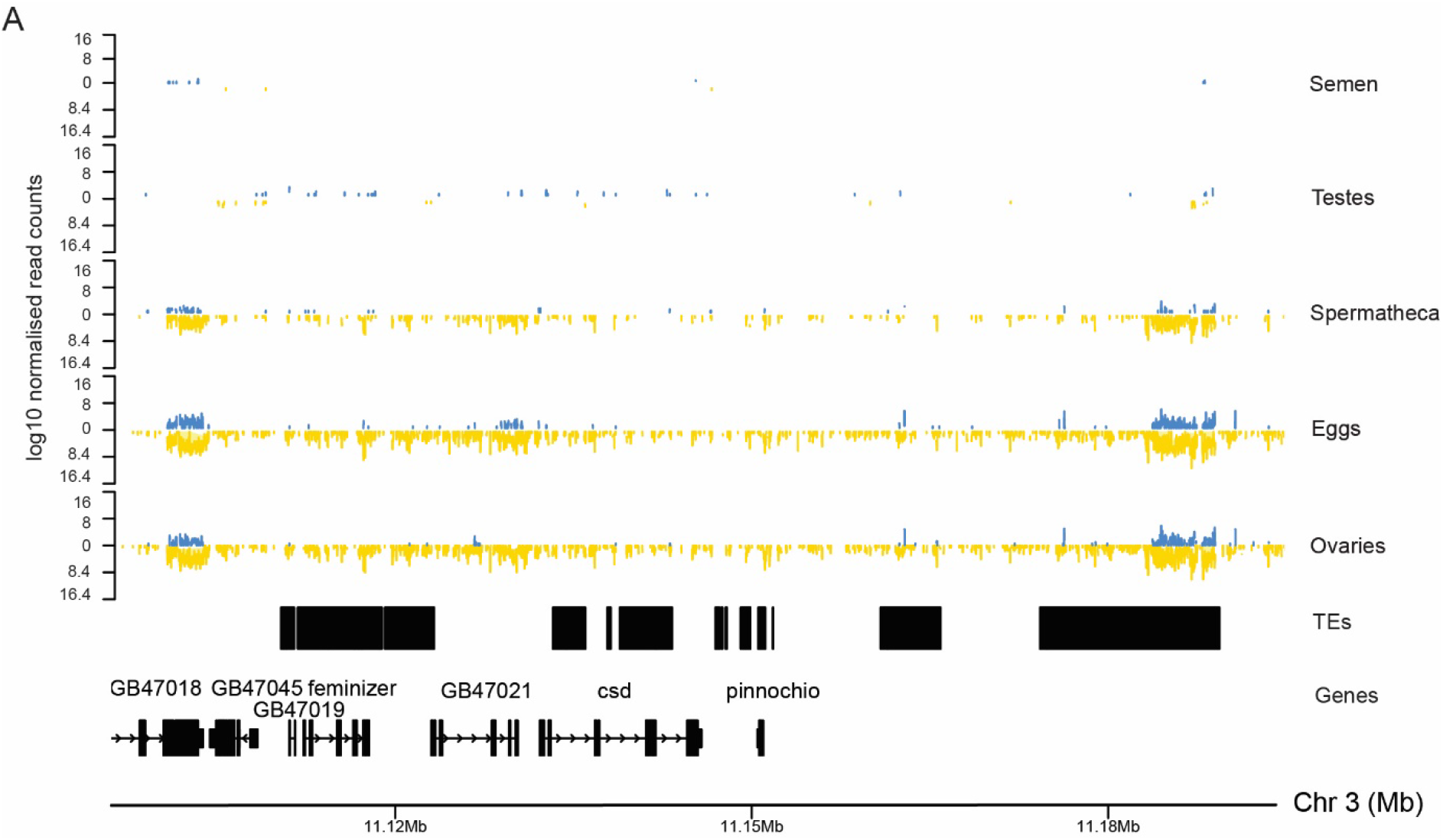
Normalised counts of piRNAs at the csd locus (Cluster 20).

## Discussion

Our pairwise comparisons of reproductive tissues of the honey bee has revealed strong differences in small RNA abundance, species and the genomic features that they target. Strikingly, these differences are strongly indicative that small RNAs function during gametogenesis, in the production, maturation and storage of semen, and in sex determination. Unique small RNA profiles are seen in fertilized and unfertilized eggs, and these differ from those seen in the spermathecal fluid. While our study was by its nature an exploratory survey rather than a test of specific hypotheses, it has revealed piRNAs and miRNAs are present in ovaries and semen and that these target genes involved in DNA regulation and developmental processes. tRNA fragments are abundant in semen and have a similar profile to those seen in the semen of other animals. We conclude that small RNAs are likely to play an integral role in honey bee gametogenesis and reproduction and provide a plausible mechanism for parent-of origin-effects on gene expression and reproductive physiology. Below we discuss the potential function of the various small RNA classes in different tissues.

### Small RNAs as potential mediators of paternal effects

tRFs made up around 60% of total small RNAs in semen, and although less abundant in spermatheca, the proportional abundance of tRF species was similar, suggesting that a large proportion of the tRFs present in the queen’s spermatheca are transferred there via spermatozoa or seminal fluid. The low frequency of tRFs in honey bee testis suggests that honey bee semen acquires tRFs from epididymosomes during transit through the epididymis, a process that coincides with spermatozoa acquiring their fertilising ability and motility. This process also occurs in mammals (Sharma et al. 2018, 2016) where seminal tRFs, particularly tRF^GlyGCC^, are acquired during epididymal transit. In mammals, tRFs are involved in the repression of embryonically expressed genes in developing offspring (Sharma et al. 2018, 2016; Schorn et al. 2017; Conine et al. 2018; Cropley et al. 2016). The high abundance of tRF^GlyGCC^ in honey bee semen highlights a striking evolutionary conservation and suggests that tRFs have a role in mediating paternal effects in honey bees and potentially in other insects. For example, Liberti and colleagues (Liberti et al. 2019) suggest that factors in seminal fluid cause queens to cease mating flights, a potential point of sexual conflict. Queens should prefer many matings to increase offspring diversity (Oldroyd and Fewell 2007) and to reduce worker-worker conflicts (Ratnieks 2015), whereas drones can maximise their probability of fathering offspring if their partner queen’s mating flights cease after his mating.

The structure of a honey bee society is predicated by the fact that a single diploid queen mates with 20 or more haploid males to produce a colony of half-sisters. It is well established that not all sub-families are equal; some are more likely to be reared as queens (Withrow and Tarpy 2018), and workers of some rare subfamilies are much more likely to become reproductively active than average (Châline et al. 2002; Oldroyd et al. 1994; Montague and Oldroyd 1998). Further, offspring gene expression and phenotype are influenced by paternity in reciprocal crosses between honey bee subspecies (reviewed in Oldroyd and Yagound (Oldroyd and Yagound 2021)).

Our work implicates seminal tRFs as a potential mechanism by which these effects may be mediated by affecting gene expression during the early development of the embryo. Further, the presence of extracellular vesicles in semen shows a potential route for transmission for tRFs from father to daughter. Nonetheless, it is difficult to see how tRFs produced by males could be transferred to eggs at an abundance that is sufficient to reliably influence embryonic development. In this context it is important to note that queens use stored sperm from the spermatheca to fertilise eggs over a lifespan that can exceed four years. This period of storage provides the opportunity for queens to ameliorate any paternal effects mediated by semen. We conclude that the function of paternal-origin tRFs in semen and spermatheca is far from clear.

Most novel miRNAs were identified in pupal testes and spermatheca. miR-210 was highly upregulated in testes, semen and spermatheca relative to ovaries and eggs. miR-210 is one of the most well-studied miRNAs across a range of species and is a master regulator of gene networks controlling neuronal development, circadian rhythm and photoreception (Cusumano et al. 2018). In *D. melanogaster*, miR-210 overexpression during development causes visual defects, however miR-210 expression is also required for photoreceptor maintenance and survival via regulating acetyl coenzyme A synthetase expression (Weigelt et al. 2019; Lyu et al. 2021). As strictly regulated expression of miR-210 is required for vision in *D. melanogaster*, miR-210 of paternal origin may be required for appropriate vision development in honey bees. miR-210 is also a candidate molecule to signal queens to cease mating flights after a successful mating (Liberti et al. 2019). Similarly, miR-34 is upregulated in testes, semen and spermatheca relative to eggs. Interestingly, miR-34 is maternally inherited in Drosophila and regulates synaptogenesis (McNeill et al. 2020) and neuronal differentiation during embryogenesis (Soni et al. 2013).

### Small RNAs may be maternally deposited in honey bees

In *D. melanogaster* piRNAs are maternally deposited in eggs, and this contribution is required for TE repression in the early embryonic development of offspring (Brennecke et al. 2008; Akkouche et al. 2013; Chambeyron et al. 2008). The high abundance of piRNAs that mapped to TEs in both egg and ovary samples suggests that the same is true in honey bees. During oogenesis in the honey bee ovary, intercellular cytoplasmic bridges allow cytoplasmic contents to be directly transferred from ovarian nurse cells into the oocyte in a process known as “nurse cell dumping” (Cavaliere et al. 1998). This suggests that, as is the case in *Drosophila*, piRNAs are transferred from honey bee nurse cells into the oocyte, priming it for TE repression upon zygotic genome activation. In addition to their role in TE repression, we found antisense gene-associated piRNAs in all reproductive tissues, indicating that some piRNAs function in gene regulation, rather than TE suppression, as has been shown in other species (Gou et al. 2014; Zhang et al. 2015). Interestingly, we found that glutamate receptor signalling genes are enriched as targets of piRNAs in maternal tissues, whereas anatomical development genes are enriched as targets of piRNAs in semen and testes. This is suggestive that piRNAs are involved in regulating developmental processes post-fertilisation as has been shown in *D. melanogaster* (Lempradl et al. 2021).

Eggs of mated queens, (both fertilised and non-fertilised) showed different small RNA profiles to virgin queens. Queens have control over egg fertilisation, and can therefore control the sex of their offspring (Ratnieks and Keller 1998). It is possible that virgin queens add more spermathecal fluid to their eggs than mated queens, perhaps as an attempt to compensate for a lack of sperm in their spermatheca, and that this alters the small RNAs present in the eggs of mated and virgin queens. In order to study this rigorously, freshly laid eggs (<2 hr) would need to be collected to detect differences before development begins. We note that in order to induce oviposition in virgin queens it is necessary to subject them to two rounds of CO2-induced narcosis (Mackensen 1947) which is a potential confounding factor (Manfredini et al. 2015). However, even if the changes in small RNA profiles that we observed in the eggs of virgin queens are caused by the CO2 treatment of the queens, it still indicates a maternal effect and communication of environmental factors between generations via small RNA molecules.

### Control of transposable elements

We have identified evidence for ping-pong amplification of piRNAs at specific piRNA clusters and TEs (see below). However, the vast majority of piRNAs in honeybee reproductive tissues arise via primary biogenesis. The overall paucity of ping-pong signatures and the high proportion of sense-TE mapping reads at many piRNA clusters, including the highly active piRNA cluster 83, indicates that most piRNAs are derived from now-inactive TE remnants, as is the case in *Drosophila* (Senti and Brennecke 2010). While most piRNAs show no ping-pong signature, piRNAs targeting annotated TEs in female reproductive tissues mapped mostly to the PiggyBac and Mariner TE families, as shown previously (Wang et al. 2017), and had much stronger ping-pong signatures than piRNAs that mapped to unclassified TEs (probably inactive remnants) (Elsik et al. 2014). This suggests that PiggyBac and Mariner TEs are transcribed in female reproductive tissues, with potential for transposition, therefore requiring robust piRNA-mediated silencing that involves ping-pong amplification. In contrast, in semen the retrotransposon LINE R2 is the only TE highly targeted by piRNAs, suggesting that it is transcriptionally active. Interestingly, LINE-1 retrotransposons produce functional reverse transcriptase in murine spermatozoa (Giordano et al. 2000), and have even been suggested as a mechanism by which extrachromosomal RNA carried by sperm could influence gene expression in progeny, perhaps by negatively regulating miRNA biogenesis (Spadafora 2017). As such, paternally inherited piRNAs that silence LINE-1 reverse transcription may permit biogenesis of certain miRNAs. Our data suggests that the presence of active LINE retrotransposons, and thus their reverse transcriptase, is highly conserved in spermatozoa, suggesting that drones may be able to influence embryonic development of their daughters via inhibition of LINE-1 activity.

### A role for piRNAs in sex determination?

The unidirectional piRNA cluster 20 encompasses two key genes involved in the sex determination pathway: *csd* and *feminizer*. This cluster includes abundant piRNAs that align antisense to *csd*. We speculate that piRNAs transcribed from this locus may regulate the expression or splicing of genes involved in sex determination as has been demonstrated in the silk moth *Bombyx mori*, where a single piRNA transcribed from *feminizer* silences a gene that controls masculinization in male embryos (Kiuchi et al. 2014) Involvement of piRNAs in sex determination is also known in *C. elegans*, where an X-chromosome derived piRNA silences a regulator of X-chromosome dosage compensation and sex determination (Tang et al. 2018). In fertilized honeybee embryos, *csd* heterozygosity leads to an active CSD protein that splices *feminizer* transcripts. The spliced Fem protein is active and in turn splices *doublesex* transcripts, leading to female development (Biewer et al. 2015). Hemizygosity for *csd* leads to normal male development whereas homozygosity leads to diploid males that are eaten by workers early in embryonic development (Woyke 1963). RNAi knockdown of *feminizer* and/or *csd* results in female to male phenotypic sex reversion (Beye et al. 2003) as does knock out of *feminizer* and/or *doublesex* (McAfee et al. 2019). It is not clear how heterozygosity at the *csd* locus results in fertile diploid females: one suggestion is that *csd* alleles form an active heterodimer (Beye 2004). Our finding that many piRNAs present in female reproductive tissues map antisense to the *csd* locus introduces another possibility; that piRNAs transcribed from one *csd* allele may affect expression or splicing of the second *csd* allele. In such a case a high level of sequence similarity between two *csd* alleles may be compensated for by piRNA-mediated silencing that results in female development despite a level of genetic homozygosity that would normally produce a male.

## Conclusions

The germline transmission of small RNAs in honey bees has not previously been studied. This study is the first to conduct pairwise comparisons of small RNA expression between reproductive tissues of maternal and paternal origin. We find evidence that, as with mammals, nematodes, and *Drosophila*, small RNAs are intimately involved in the regulation of gametogenesis and embryogenesis in the honey bee and provide a plausible but tentative pathway for parental manipulation of gene expression in offspring. Future studies will seek to experimentally validate the role of small RNAs at key loci such as the sex-determination locus (*csd* and *feminizer*) and to uncover the role of seminal tRFs. Studies such as these are essential to illuminating our understanding of how epigenetic mechanisms evolve to promote individual fitness.

## Methods

### Sample Collection

#### Semen

In February 2015 mature drones were sampled randomly from 3 colonies. Semen from 10 drones per colony was collected into sterile glass insemination tips using standard procedures used during honey bee artificial insemination (Harbo 1986). Semen from each sample was expelled directly into 0.5 ml Trizol and the tube vigorously flicked to disperse the semen in the reagent. The preparation was then frozen at −70 until RNA extraction.

#### Eggs from unmated queens

In January 2015 we established four nucleus colonies and introduced one queen pupa to each using standard methods {Harbo, 1986, Propagation and instrumental insemination}. The sister queens were prevented from leaving the nucleus colonies via a ‘queen excluder’ grid placed over the entrance which was sufficiently large to allow the passage of workers, but too small to allow the passage of queens. When the queens were 7 days old we narcotized them with carbon dioxide for 10 minutes, once a day for two days, to induce oviposition, even though they had not mated (Mackensen 1947). Once the queens hard started laying we provided drone comb and worker comb to the colonies. Over several weeks we collected eggs 0-24 hours old into 0.5 ml Trizol from drone comb and worker comb. Eggs disintegrated on contact with the Trizol. Each sample consisted of 59-200 eggs. Samples were frozen at −70 C after collection.

#### Eggs from mated queens

Eggs were collected in September 2015 from three colonies headed by naturally mated queens of unknown age. Eggs were collected separately from drone and worker cells into 0.5 ml Trizol as above.

#### Spermathecae

In March 2016 10 naturally-mated laying queens were removed from their colonies and taken to the laboratory. Queens were frozen, and the ovaries and spermatheca were removed by dissection as described in Dade (1977) (Dade 1977). Tissue was placed separately in 0.5 ml Trizol and immediately frozen.

#### Testes

In August 2016, a section of brood comb containing newly capped drone larvae was collected from a single colony and taken into the laboratory. White eyed drone pupae were extracted from their cells and placed in petri dishes lined with filter paper soaked in 12% glycerol. Pupae were incubated at 34.5 C, 50% relative humidity, and staged to collect developing testes at three time points. On day 1 (24 hours post-incubation), pink-eyed pupae were collected and testes removed from the abdomen by piercing the cuticle with forceps and aspirating testes tissue using a P1000 pipette tip that had been widened by removing the end of the tip with sterile scalpel. Red-eyed pupae were collected on day 3 (72 hours post-incubation) and brown-eyed pupae on day 6 (144 hours post-incubation) and testes removed as described. Samples were frozen immediately on dry ice and stored at −80 C prior to RNA extraction. Two individual drone pupae samples were used to generate two replicate small RNA libraries for each timepoint.

### RNA extractions

Frozen tubes containing the sample and 500 μl Trizol were thawed on ice. The sample was then homogenized with a RNAse free pellet pestle and RNA extracted using Trizol according to the manufacturer’s instructions (Life Technologies).

### Transmission electron microscopy

For Extracellular Vesicles’ imaged using transmission electron microscopy (TEM), samples were diluted 1:200 in PBS and applied to formvar-coated grids with heavy carbon coating after fixation in glutaraldehyde (2% v/w in H2O) for 30 mins, room temperature, or overnight at 4oC. Samples were visualized by negative staining with uranyl acetate (2% w/v in H2O), with images captured using a Joel JEM-2100 electron microscope.

### Nanoparticle tracking analysis of isolated exosomes

Extracellular Vesicles’ were analyzed for size and concentration via the Zetaview instrument model Basic-PMX120 installed with a 405 nm laser diode (Particle Metrix). Vesicles were prepared with a 1:20,000 dilution in dPBS to an average 149 particles per frame. For each measurement, “high” (60fps) number of frames were captured for 11 positions; Camera sensitivity was 80; Shutter was 100; After capture, analysis was performed by ZetaView Software version 8.05.12 SP1.

### Small RNA Library prep and mapping

Small RNA libraries were prepared according to the manufacturer’s instructions (Illumina TruSeq Small RNA Library Prep Kit, Part # 15004197 Rev. G) unless stated otherwise. Input amount for library preparation was 1.5 μg of total RNA, quantified using a NanoDrop Spectrophotometer. PCR amplification was performed for 15 instead of 11 cycles. The amplified cDNA construct was purified on 6% TBE polyacrylamide gels. Gel bands between 145 bp and 160 bp in size, which correspond to adapter ligated 22 nt and 30 nt RNA fragments, were excised and purified. The final library was concentrated by ethanol precipitation and the pellet was resuspended in 10 μl 10 mM Tris-HCI, pH 8.5. Libraries were sequenced at the Australian Genome Research Facility using the Illumina HiSeq-Single Read (50/100) chemistry.

Small RNA libraries were mapped to the Amel4.5 genome using CLC Genomics (Qiagen). Biotype annotations were assigned using the Annotate and Merge tool.

### Novel miRNA identification

Novel miRNA prediction tool mirDeep2 (Friedländer et al. 2012) was used to predict novel miRNAs in each tissue (biological replicates pooled). CLC Genomics (Qiagen) was used to manually confirm expression of each predicted miRNA: those for which expression could not be confirmed (expression <5 in all tissues) were removed.

### Validation of miRNAs by stem–loop RT–PCR

For the validation of miRNAs by stem–loop RT–PCR independent samples were collected in September 2017. Total RNA of pink, red and brown testis, semen, eggs, ovaries and spermathecae was extracted, procedure for tissue collection and RNA extraction as described above. The protocol for miRNA validation by stem-loop RT-PCR was adapted from Ashby et al (2015) (Ashby et al. 2016) and Varkonyi-Gasic et al (2007) (Varkonyi-Gasic et al. 2007). Primers for stem-loop RT-PCR were designed with a miRNA primer design tool (Astridresearch: http://genomics.dote.hu:8080/mirnadesigntool/processor), using the base stack melting temp primer (Czimmerer et al. 2013) (Supplemental Table S8).

For Reverse Transcription in a final volume of 20 ul 50 ng of total RNA was combined with 50 nM of the StemLoop RT primer, 0.25 mM dNTP mix and nuclease free water to 13.65 ul. The mixture was heated to 65°C for 5 minutes followed by a 2-minute incubation on ice. 6.35 ul of the enzyme mix consisting of 1x First First-Strand buffer, 10 mM DTT, 4 U RNaseOUT, 50 U SuperScript III Reverse Transcriptase (Life Technologies, Australia) was added and incubated in a Biorad Mycyler thermalcycler under following conditions: 16 °C for 30 minutes, 42 °C for 30 minutes, heat inactivated at 85 °C for 5 minutes and cooled to 4 °C. RT-PCR was performed as described by Ashby et al (2015). For a 15 ul reaction 1 ul of undiluted cDNA was combined with 1x FastStart Universal Probe Master Mix with Rox (Roche Diagnostics, Australia), 50 nm of the forward and universal reverse primer, 10 nm of Universal Probe #21 (Roche Diagnostics, Australia). Cycling conditions were 95 °C for 5 minutes, 40 cyles of 95 °C for 10 seconds, 56 °C for 30 seconds, 72 °C for 10 seconds on a Applied Biosystems FAST 7500 real-time PCR machine using the normal ramp reaction mode.

### piRNA Analysis

Prior to cluster identification, unmapped reads and reads that were annotated as other known RNAs (such as ribosomal and tRNAs, but excluding protein coding RNA) were removed. A custom script (CREST) was generated to identify genomic regions containing a high density of piRNA reads. After clusters were identified, those which contained less than 50% of reads of the appropriate size (26-31nt) were filtered out. proTRAC v2.4.2 (Rosenkranz and Zischler 2012) and piClust (Jung et al. 2014) were run using the same dataset but excluding reads less than 26nt or greater than 31nt in length to ensure consistency across packages. Default parameters were used for proTRAC v2.4.2 and piClust was run using parameters; eps of 10,000, minread of 35 and a cut score of 2. Tissue-specific clusters which were either overlapping or within 1kb of each other were merged and clusters less than 200 bp in length were removed. Cluster strandedness was determined by calculating the proportion of collapsed reads mapping to either strand. If one strand mapped more than double the number of piRNAs relative to the other strand, across all tissues, it was assigned as either sense or antisense. If there were fewer than double the number of reads (collapsed) on either strand, across all tissues, the cluster was assigned as a dual strand/bidirectional cluster. We identified enrichment of 10 nucleotide overlaps using the tool PPmeter (v 0.4) (Jehn et al. 2018) and phased piRNA signatures were identified by calculating the 3’-5’ distances between putative piRNAs. piRNAs were mapped to the Amel 4.5 TE dataset (Elsik et al. 2014), TE annotations are divided into TE class and TE superfamily as many repetitive elements could not be assigned to a superfamily (ie. *Mariner, Copia etc*) but were assigned to a class (ie *LARD, TRIM etc*).

### Target prediction and gene ontology analysis

3’UTR sequences were obtained from OGS v3.2 (Beebase). Two miRNA target prediction algorithms, miRanda (Betel et al. 2010) and RNAhybrid (Kruger and Rehmsmeier 2006) were used to identify miRNA 3’UTR-gene targets. Only miRNA-target interactions predicted by both algorithms were used for GO analysis. The targets were matched against each list of differentially expressed miRNA, for each pairwise tissue comparison, and the genes searched using the graphical gene set enrichment tool ShinyGO v 0.61 to obtain enriched biological process, cellular component and molecular function terms, KEGG pathways and promoter motifs {Ge, 2020 #55}. To determine the cellular processes that piRNAs may regulate in each tissue type we sought to identify genes and TEs that are differentially targeted for each pairwise tissue comparison. Genes and TEs which had significantly more putative piRNAs mapping antisense to them in one tissue relative to another were assigned as differentially targeted. The Featurecounts command, as part of the Rsubread package (v. 1.22.2), was used to counts reads mapping to TEs and genes (including introns, exons, 3’UTR and 5’UTR). Intergenic reads were calculated by subtracting reads mapping to TEs and/or gene elements from the putative piRNA count (after length filtering).

## Data Access

All raw and processed sequencing data generated in this study have been submitted to the NCBI Gene Expression Omnibus (GEO; https://www.ncbi.nlm.nih.gov/geo/) under accession number GSE182720.

## Acknowledgments

This work was supported by Australian Research Council grant DP150100151 to BPO and AA. AA is supported by the Australian Research Council (DE140100199 and FT180100653). The authors thank Dr Ros Gloag for helpful discussions about the *csd* locus.

